# Modeling the Mechanism of CLN025 Beta-Hairpin Formation

**DOI:** 10.1101/145185

**Authors:** Keri A. McKiernan, Brooke E. Husic, Vijay S. Pande

## Abstract

Beta-hairpins are a substructure found in proteins that can lend insight into more complex systems. Furthermore, the folding of beta-hairpins is a valuable test case for benchmarking experimental and theoretical methods. Here, we simulate the folding of CLN025, a miniprotein with a beta-hairpin structure, at its experimental melting temperature using a range of state-of-the-art protein force fields. We construct Markov state models in order to examine the thermodynamics, kinetics, mechanism, and rate-determining step of folding. Mechanistically, we find the folding process is rate-limited by the formation of the turn region hydrogen bonds, which occurs following the downhill hydrophobic collapse of the extended denatured protein. These results are presented in the context of established and contradictory theories of the beta-hairpin folding process. Furthermore, our analysis suggests that the AMBER-FB15 force field, at this temperature, best describes the characteristics of the full experimental CLN025 conformational ensemble, while the AMBER ff99SB-ILDN and CHARMM22* force fields display a tendency to overstabilize the native state.

## I. INTRODUCTION

Molecular dynamics (MD) simulations employ a potential energy function, referred to as a force field, in order to sample the free energy landscapes of biomolecular systems. Due to the intractable complexity of biological systems, the force field is commonly of an approximate classical form, and is fit using quantum mechanical and experimental data^1,2^. As a result, the accuracy of these force fields has tracked with advances in computational hardware and methodology, as well as the increased availability of high resolution experimental data^3^. The current state-of-the-art protein force fields demonstrate high accuracy in their ability to describe the protein native state and its equilibrium behavior: these models are all able to describe ensemble averaged properties of proteins with highly populated native states within experimental error^4^. Of greater discrepancy is their description of the denatured state ensemble. As such, one of the major frontiers in protein force field development is the accurate description of proteins away from equilibrium.

Protein folding is a strong validation test of a protein force field^5,6^. This is because as the protein folds and unfolds, it samples beyond the native state. Performing protein folding simulations using multiple force fields allows for the comparison of their denatured state ensembles. Furthermore, we can make force field-agnostic conclusions from an aggregated dataset. A popular system for such a task is the ultrafast folding peptide, CLN025^7^. CLN025 is a 10-residue peptide designed based on the C-terminal fragment of Protein G (residues GLY41 to GLU56)^8^. Because the CLN025 peptide adopts a unique structure at room temperature, can be crystallized, and possesses a funnel-like free energy surface, Honda *et al.* ^7^ argue that it should be considered a protein. We will thus refer to CLN025 as a miniprotein, which captures both its small size and exceptionally stable behavior.

CLN025 folds within timescales accessible to computation into a highly stable beta-hairpin. At room temperature, the native state is almost exclusively populated^7^. As temperature increases, the denatured state population increases. Experiments probing relaxation kinetics over a range of temperatures have shown that there is a critical break in the folding mechanism of this miniprotein at 308 K^9^. Above this temperature, folding can no longer be described using a two-state model. Because the experimental description of this system is both detailed and nontrivial at high temperature, we have benchmarked a set of popular protein force fields in their ability to describe the conformational dynamics of CLN025 at its experimental melting temperature of 340 K^7^ (see Sec. II).

The folding of CLN025 is of additional interest due to its beta-hairpin structure. Many theoretical^10–35^ and experimental^9,36–47^ studies have presented conflicting mechanisms and rate-limiting steps for beta-hairpin formation. Mechanistic studies have somewhat converged on two leading mechanisms that were developed to explain the folding of the C-terminal fragment of Protein G. The first, suggested by Muñoz *et al.* ^36,37^ to explain relaxation kinetics observed from T-jump experiments, proposes that the hairpin turn forms first from the extended state. The beta-sheet then “zips” from the turn to the terminus via the formation of a series of cross-strand hydrogen bonds; in so doing, the structure becomes collapsed. The second mechanism, formulated by Pande and Rokhsar ^12^ and by Dinner, Lazaridis, and Karplus ^11^, proposes that hydrophobic collapse occurs first and the turn is formed from the collapsed structure. The mechanism of Pande and Rokhsar ^12^ includes the same “zipping” of hydrogen bonds from the turn to the terminus, whereas the mechanism of Dinner, Lazaridis, and Karplus ^11^ proposes formation of hydrogen bonds starting near the middle of the beta sheets and propagating outward in both directions.

Both experimental and theoretical studies have corroborated the turn zipper model^22–28,36–39^ or the hydrophobic collapse model^9–22,43–45^ for the Protein G fragment as well as other beta-hairpin containing systems^22–24,31,35,39,42–47^ including CLN025^9^ and its predecessor chignolin^21,28,32,34^. Additionally, some have proposed cooperative mechanisms where turn formation and hydrophobic collapse occur simultaneously^30,31,40,41^ or alternative mechanisms^27,34^. There is also much disagreement regarding the rate-limiting step of beta hairpin formation. Muñoz *et al.* ^36,37^ and subsequent studies^28,31,39^ found that the formation of the turn from the extended state determines the formation rate. For the hydrophobic collapse mechanism, some groups argued that the positioning of hydrophobic groups so that the cluster can form determines the rate^12,17,43^ while others claimed an interconversion between compact conformations is the rate-limiting step^9,11,19,43–45^. Additional studies implicated the formation of interstrand contacts and hydrogen bonds^13,32,34^.

Notably, interrogations of the chignolin and CLN025 folding mechanisms have mirrored the larger beta-hairpin folding community in producing irreconcilably different results. MD simulations of chignolin performed by Suenaga *et al.* ^21^ showed a folding mechanism in which hydrogen collapse precedes formation of the proper turn structure, while a different MD study of chignolin by Harada and Kitao ^28^ showed agreement with the turn zipper mechanism. In their 2011 seminal protein folding study, Lindorff-Larsen *et al.* ^33^ simulated CLN025 but did not comment on a specific mechanism. In 2012, Davis *et al.* ^9^ performed the first experimental interrogation of the CLN025 folding mechanism and reported significantly faster timescales for beta sheet and hydrophobic collapse than for the turn formation, suggesting the hydrophobic collapse occurs first.

In light of the vast literature surrounding beta-hairpin formation and the recent experimental results for CLN025 folding^9^, we use our aggregated MD dataset to facilitate the understanding of beta-hairpin formation. We first enumerate the force fields studied and discuss the Markov state model (MSM) framework used to analyze our MD datasets. Next, we examine the thermodynamics and kinetics of folding for the three force fields investigated and note that only the AMBER-FB15 model^4^ exhibits melting behavior at the simulation temperature. Lastly, we analyze the three MD datasets simultaneously to interrogate the mechanism and rate-determining process of CLN025 folding. Through this analysis we find that the CLN025 folding mechanism comprises a downhill hydrophobic collapse followed by the slower formation of the hairpin turn over a barrier. The order of these conformational changes is consistent with the recent experimental study of CLN025^9^.

## II. METHODS

### A. Simulations

The force field combinations used in this study are:

(a) CHARMM22*^48^/mTIP3P^49^

(b) AMBER ff99SB-ILDN^50^/TIP3P^51^

(c) AMBER-FB15^4^/TIP3P-FB^52^

The CHARMM22* and AMBER ff99SB-ILDN parameter sets were developed by Piana, Lindorff-Larsen, and Shaw ^48^ and Lindorff-Larsen *et al.* ^50^, respectively, as augmentations to previous generations of CHARMM and AMBER parameter sets. The AMBER-FB15 parameter set, developed by Wang *et al.* ^4^, was built via a complete refitting of the bonded parameters of the AMBER ff99SB force field^53^ with training data taken from RI-MP2 calculations using augmented triple-zeta and larger basis sets^54^. Notably, the training set contained complete backbone and side chain dihedral scans for all (capped) amino acids. During force field validation, it was found that parameter optimization yielded improved melting curves for both CLN025 and Ac-(AAQQAA)_3_-NH_2_. We suspect that the improved thermal dependence could be attributed to improved description of the dihedral barrier heights ^55–57^. Note that each protein force field was simulated using its corresponding water force field. The dataset for model (a) was obtained from D.E. Shaw research, and was simulated at 340 K as described in their fast folding protein study^33^. The dataset comprises one (continuous) 106 *μ*s simulation divided into 54 segments for file transfer purposes.

The datasets for models (b) and (c) were generated via the distributed computing platform Folding@home^58^. To prepare these simulations, the CLN025 crystal structure was first solvated (3961 water molecules) and neutralized (2 Na^+^ ions). The system was denatured via simulation at 600 K until fully extended, using the AMBER ff99SB-ILDN parameterization. For model (c), this configuration was then ported to the AMBER-FB15 and TIP3P-FB force fields. The configuration was then equilibrated for 1 ns at 340 K. After equilibration, 100 instances of this positional configuration were written as initial conditions for a unique Folding@home simulation, each with a unique velocity distribution. This architecture allows us to sample many instances of folding from the extended state, and hence gather robust statistics regarding the folding process. The Folding@home simulation dataset for AMBER ff99SB-ILDN/TIP3P contains 141 independent simulations up to 1 *μ*s in length for a total of 103 *μ*s and the dataset for AMBER-FB15/TIP3P-FB contains 144 independent simulations up to 1 *μ*s in length for a total of 101 *μ*s. The OpenMM^59^ script used to convert the denatured system to the set of Folding@home initial states as a function of input protein and water force fields is provided in the supplementary materials.

### B. Markov state models

Whereas specialized hardware is typically used to generate one or several ultralong MD simulations, simulations performed on distributed computing platforms such as Folding@home produce datasets consisting of many short trajectories. The use of MSMs was a crucial advance in the analysis of such datasets^60–63^. To construct a MSM, each frame of each trajectory is assigned to a discrete state. The model comprises the populations of and conditional pairwise transition probabilities between states, which provide thermodynamic and kinetic information, respectively. Since separate trajectories will feature common states, the trajectories can be threaded together through this framework and pathways between states can be determined even if the pathway is not contained in a single trajectory.

The Markov assumption underlying MSMs is that the states are “memoryless”: the probability of the system transitioning from one state to another is independent of the previous states the system was in. In order to satisfy the memorylessness requirement, transitions within states must occur much faster than transitions among states. The choice of which collective variables to use to determine these states is an area of active research^64^. Describing the trajectories using time-structure based independent component analysis (tICA) allows us to analyze the MD dataset in terms of its slow dynamical processes^65,66^. Each component of the tICA transformation, or “tIC”, serves as a reaction coordinate for the system^67^. For a protein folding dataset, the first tIC is expected to correspond to the folding process and can thus be used as a reaction coordinate for folding. The MSM is then created from trajectories that are represented by their progress along the tICs by creating microstates that group kinetically similar conformations. This representation is therefore expected to produce states that satisfy the Markov property^65^. Verifying adherence to the Markov property is extensively discussed elsewhere^63^ and model validation for the models used here is presented in the supplementary materials.

#### 1. Projected MSMs

In this report we use two types of MSMs for analysis. We refer to the first type as a “projected” MSM which is projected from a “baseline” MSM. The baseline MSM is created from the CHARMM22* dataset. First, the dataset was featurized into the sines and cosines of the *α* dihedral angles (i.e. the dihedrals along the *α*-carbon backbone) and transformed using the tICA algorithm^65,66^ with a lag time of 128 ns and the kinetic mapping^68^ weighting scheme. All 14 components of the tICA solution were retained and clustered into 704 microstates using mini-batch *k*-means. A Markov state model (MSM) was constructed on the entire dataset with a MSM lag time of 50 ns based on the lag time reported by Beauchamp *et al.* ^69^ for the same dataset. Details regarding the optimization of these parameters and model validation are provided in the supplementary material.

To create the projected MSMs for the AMBER datasets, in accordance with the baseline MSM, the AMBER datasets are first featurized into the sines and cosines of the *α* dihedral angles. Then, to perform the projection, the baseline tICA and clustering models prefit on the CHARMM22* dataset are used to predict (1) the location of each AMBER frame along the tICs and (2) the microstate assignment of that frame. Each projected MSM was built from these predicted assignments. By using projected MSMs, the tICs and microstates are the same for all models. Using this framework, state populations and pairwise transition probabilities can be directly compared.^70^ This type of model is used in Sec. III A and Sec. III B below.

#### 2. Subprocess MSMs

The second type of MSM used in this work is a “subprocess-specific” MSM. In order to estimate the rate of a specific dynamical process, only the degrees of freedom involved in that process are used to construct the model. This method is developed as an analogy to Tjump experiments so that specific relaxation processes can be isolated. In this work, we hand-select features that correspond to the hydrophobic collapse and turn formation processes of CLN025 folding in order to separately analyze their timescales and reaction coordinates. This type of model is used and further described in Sec. III C and in the supplementary material.

#### 3. Model statistics

Bootstrapped Markov state model construction allows for the comparison and visualization of statistics. For each model and MSM variety, 100 boostrapped MSMs were created by randomly selecting trajectories from the relevant dataset with replacement. Error bars along reaction coordinates are represented as the interval between the 5^th^ and 95^th^ percentiles for free energy values calculated from each boostrapped MSM. All bootstrapped MSMs used in this work are provided as supplementary files and instructions for loading them are provided in the supplementary material.

## III. RESULTS

First, we describe folding from a global perspective and compare the thermodynamics and kinetics for each force field. Then, we inspect the mechanism of beta-hairpin formation for each dataset in the context of a set of influential theoretical and experimental studies. Last, we examine the rate-determining step of the folding process.

### A. Thermodynamics and kinetics of folding

In order to analyze folding of CLN025, we first constructed a baseline MSM for the CHARMM22* dataset. The same features, tICA model, and states were then used to derive a unique MSM transition matrix for the AMBER ff99SB-ILDN and AMBER-FB15 datasets as described in Sec. II B. By using a consistent model basis, we are able to directly compare folding of CLN025 as a function of force field^71^. This approach allows us to summarize folding along a kinetically motivated 1-dimensional reaction coordinate (see Sec. II B for more detail). This data is illustrated in Fig. 1 (top), where folding from the denatured state is represented by the movement from the right free-energy basins, labeled “denatured extended” and “denatured collapsed”, to the left basin, labeled “folded”. Additionally, in order to quantify the kinetics of folding, we computed the mean first passage time (MFPT) both to and from the folded and denatured states for each model. This is depicted in Fig. 1 (bottom) and the method is described in the supplementary material.

**FIG. 1.**
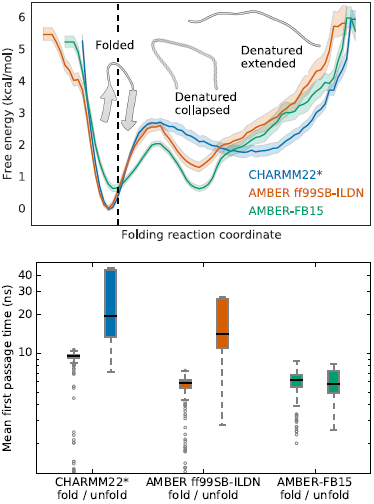
Thermodynamics and kinetics of CLN025 folding reveal differences across force fields. Top: The free energy landscape shows overstabilization of the native state for the CHARMM22* and AMBER ff99SB-ILDN force fields at the experimental melting temperature. The black dashed line indicates the location of the crystal structure (PDB ID: 5AWL) on the folding reaction coordinate (see II B). All models populate a folded state and a denatured extended state, but only the AMBER models populate a denatured collapsed state at this temperature. Shading represents the range of free energies based on 100 bootstrapped samples between the 5^th^ and 95^th^ percentiles. The folding free energy (Δ*G*folding) is computed from the change in free energy between the minima of the folded and denatured basins. In kcal per mole, Δ*G*folding is on the interval (1.56, 2.02) for CHARMM22*, (1.07, 1.51) for AMBER ff99SB-ILDN, and (-0.20, 0.18) for AMBER-FB15. Bottom: The mean first passage time for folding is approximately similar for all force fields; however, the mean first passage time for unfolding is slower than folding for the CHARMM22* and AMBER ff99SB-ILDN models at the experimental melting temperature. Only the AMBER-FB15 model shows approximately equal folding and unfolding MFPTs, which is consistent with melting behavior.

We found that all models share several notable characteristics. First, all models show that the folding process is rate limited by a small global barrier (Fig. 1, top). This demonstrates that the potential energy surfaces for CLN025 described by each of these force fields are qualitatively similar. Second, all were able to fold the extended, denatured miniprotein into a native conformation similar to the experimental crystal structure (dashed line, Fig. 1, top). The minimum of the folded basin is found at a similar location on the reaction coordinate for all models. This implies that the most stable folded conformations are also very similar. Third, the mean first passage time (MFPT) for folding was found to be on the order of 10 ns. This is evidenced by the short and comparable folding MFPTs for all models (Fig. 1, bottom).

The examined models also differ in several ways. First, their description of dynamics at the experimental melting temperature differ. During melting, the native and denatured states should be equally populated, and the folding and unfolding rates should be the same. We found that the AMBER-FB15 model displays equally deep folded and unfolded basins, well as approximately equal folding and unfolding MFPTs. This is aligned with experiment at the same temperature. In contrast, the CHARMM22* and AMBER ff99SB-ILDN models exhibit disproportionately high unfolding barriers, and unfolding MFPTs much slower than their corresponding folding MFPT. This represents overstabilization of the native state at the experimental melting temperature. Such a phenomenon is a common limitation of protein force fields^72,73^.

We expect that melting behavior would be achieved for the CHARMM22* and AMBER ff99SB-ILDN models at temperatures higher than the experimental melting temperature. The full melting curves for the CLN025 system are given for each of these force fields in Wang *et al.* ^4^. In that work, the simulated melting curves were determined via replica exchange simulations. By closely examining the dynamics at the experimental melting temperature for a subset of the most accurate and broadly used force field combinations, our results agree with and expand on the results of Wang *et al.* ^4^. In both studies, we see that the AMBER-FB15 model is 50% folded at the experimental melting temperature, while the AMBER ff99SB-ILDN and CHARMM22* models display an overly populated folded state. Additionally, the melting curves suggest that the CHARMM22* model has a melting temperature of 350 K or greater, while the AMBER ff99SB-ILDN model does not reach 50% folded over the given temperature range, and is therefore expected to be greater than 370 K. In the work of Lindorff-Larsen *et al.* ^33^, the folding temperature for CLN025 with CHARMM22* is reported to be 370 K^74^.

Our comparison of the dynamics at the experimental melting temperature shows the discrepancies of the CHARMM22* and AMBER ff99SB-ILDN dynamics with experiment at the same temperature. Since the CHARMM22* and AMBER ff99SB-ILDN models do not show melting, these models differ considerably in the structure of the denatured state ensemble. At 340 K, the AMBER ff99SB-ILDN and AMBER-FB15 models populate both an extended denatured state as well as a compact denatured state. This compact denatured state describes a hydrophobically collapsed structure. In contrast, the CHARMM22* model populates almost exclusively the denatured extended state.

In order to look more closely at the structure of the folded and denatured states, statistically representative trajectories were sampled from each MSM, and a Ramachandran plot was constructed for each pair of adjacent residues. Ramachandran plots for all residue pairs are given in the supplementary materials. We have depicted these plots for a set of two residue pairs in Fig. 2. The dihedral coordinates for the crystal structure (PDB ID: 5AWL) are denoted by the black circle. The pair TYR2–ASP3 (Fig. 2, top) is near the terminal region of CLN025, while THR6–GLY7 (Fig. 2, bottom) is located in the turn region. Each depicted pair characterizes a distinct region of the folded state. Note that in the folded state, all residue pairs across all models display structure very close to the experimental crystal structure. As expected, the folded state conformations for the terminal residue pair is located in the parallel beta-sheet region of dihedral space (*ϕ* ≈ -120*°*, *ψ* ≈ 115*°*), while the turn region residue pair is localized to the glycine region of dihedral space and is part of a sequence describing a type II*′* beta-turn (see supplementary materials for other plots in the turn sequence)^75^. These findings corroborate the tICA analysis suggesting that the folded states for all models are very similar. There are larger differences between the denatured states. The denatured state of the CHARMM22* model is diffuse and does not display a strong skew toward any particular dihedral basin. This suggests that the denatured state ensemble for this model is very extended. This is in contrast to the AMBER ff99SB-ILDN and AMBER-FB15 models, each of which shows a higher population in the major basin of the folded ensemble. This suggests that within the denatured ensemble for these models, there is a thermodynamic preference for a structure with a native-like topology.

**FIG. 2.**
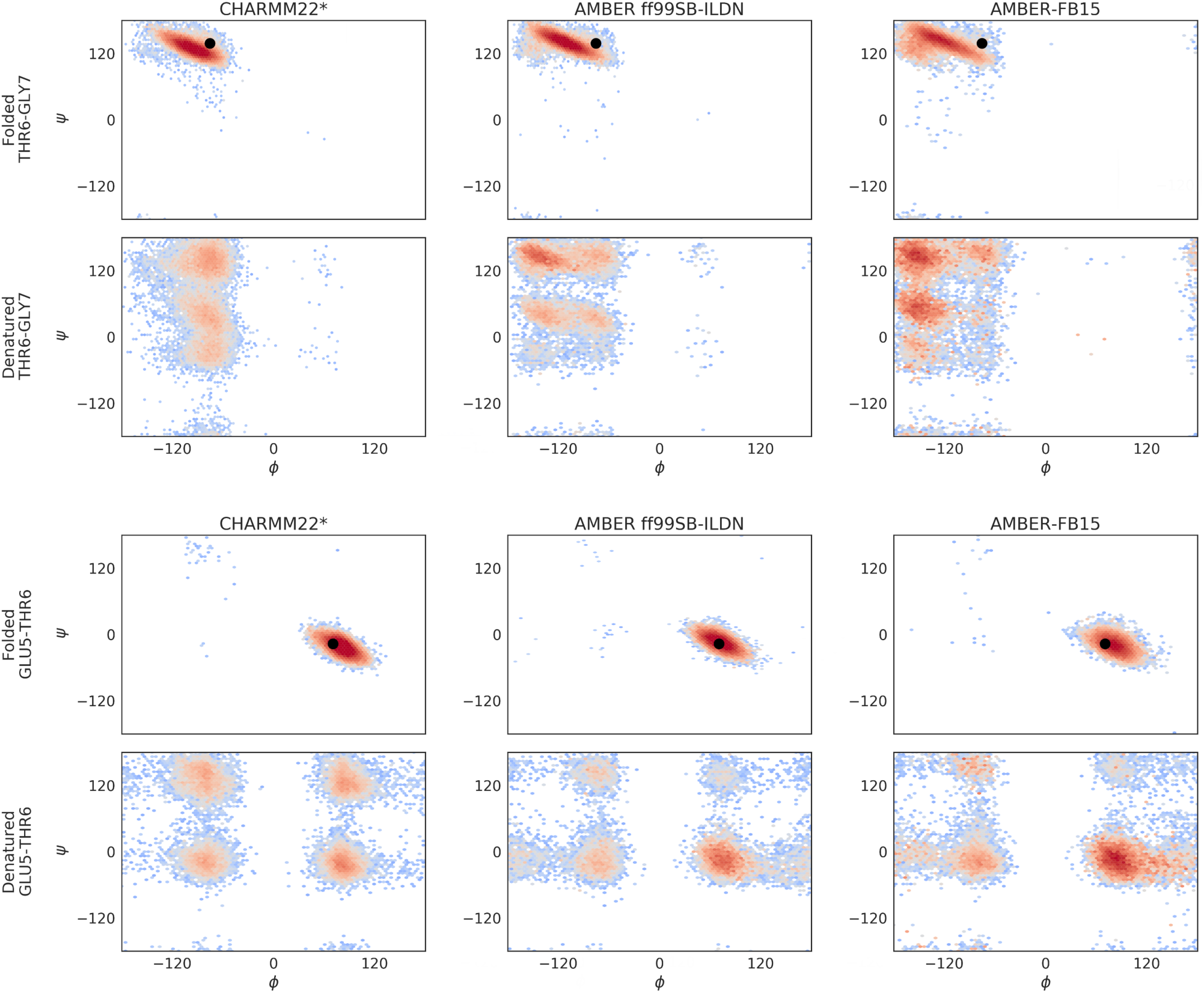
Sampling the folding MSM allows for the structural characterization of both the folded and denatured state ensembles. The depth of these basins denotes the relative thermodynamic stability of a given dihedral conformation. Here we have focused on a residue pair in the beta-sheet region of the folded structure (TYR2–ASP3), and a residue pair in the turn region (THR6– GLY7). Each column represents a force field combination. For all models, we observe that the folded state is structurally very similar to and in agreement with the experimental crystal structure (denoted by the black circle). The folded state ensemble of the beta-sheet residue pair, TYR2–ASP3, is mostly populated in the parallel beta-sheet region of dihedral space (*ϕ* ≈ -120*°*, *ψ* ≈ 115*°*). The folded state ensemble of the turn-region residue pair, THR6–GLY7, is mostly populated in the glycine region of dihedral space, and denotes part of a sequence describing a type II’ beta-turn. The models differ more with respect to the denatured state ensembles. It can also be seen that the CHARMM22* denatured states are relatively diffuse and uniform, suggesting a very extended ensemble. In contrast, the AMBER ff99SB-ILDN and AMBER-FB15 models exhibit higher population in the dihedral basins of the crystal structure, suggesting higher thermodynamic stability for conformations with a native-like topology.

### B. Mechanism of beta-hairpin formation

In 2012, Davis *et al.* ^9^ used T-jump experiments combined with infrared and fluorescence spectroscopy to empirically measure the relaxation kinetics of the turn and terminal regions of CLN025. It was found that above 308 K, folding cannot be described using a two-state model. Above this temperature,^9^ showed that the turn, beta sheet formation, and hydrophobic collapse processes occur on significantly different timescales, with a faster rate observed for beta sheet and hydrophobic cluster formation. Additionally, as temperature increases toward the melting temperature, the timescale separation increases. These results suggest a mechanism in which interactions of the terminal hydrophobic residues first cause the extended structure to collapse into a nativelike topology, after which small local rearrangements occur, forming the turn and the remaining native state contacts. While this experimental characterization describes the ordering of major conformational changes, it does not resolve the relative order of specific hydrogen bond formation in the beta sheet.

We analyzed the sequence of events in our simulation datasets by using the models created above. To assess whether the turn had formed, we tracked the existence of three hydrogen bonds characterizing the turn (purple distances, Fig. 3). To determine whether the structure had collapsed, we used a binary metric based on the radii of gyration of the two hydrophobic terminal residues (TYR1 and TYR10; dark gray residues, Fig. 3).^76^ Finally, to monitor the completion of beta sheet formation, the three terminal hydrogen bonds were monitored (dark gray distances, Fig. 3). We elaborate on these feature sets in the supplementary material.

**FIG. 3.**
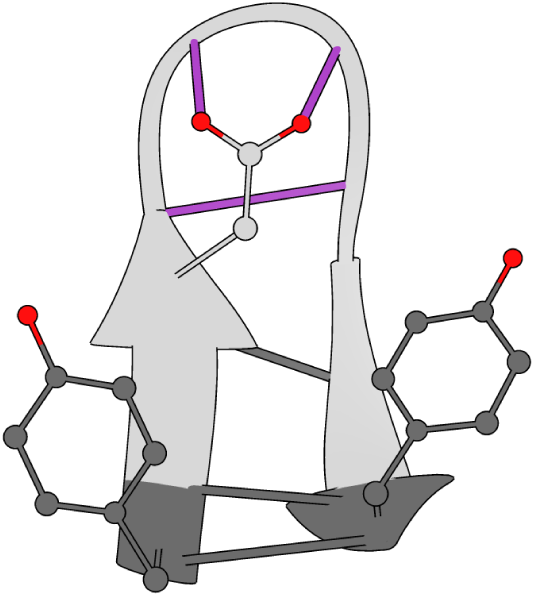
Subprocess-specific features are used to monitor the CLN025 folding mechanism. Hydrogen bonds of the native state turn, for which distances were calculated, are shown in purple. Hydrophobic residues, for which the radii of gyration were measured, are highlighted in dark gray. Distances of the native state beta sheet hydrogen bonds are also highlighted in dark gray.

To provide an intuition about the folding mechanism, Fig. 4 shows a representative trajectory for each of the three MD datasets. First, it is interesting to note that in the CHARMM22* dataset, folding occurs as a concerted mechanism: the miniprotein is either denatured extended or folded, and the turn formation and collapse occur simultaneously and quickly from the extended state. The model of CHARMM22* at this temperature does not resolve the mechanism enough to compare or contrast it with the existing theories of beta-hairpin formation. In the AMBER datasets, however, the turn and hydrophobic collapse occur gradually with instances of collapse (formation of hydrophobically collapsed structures) preceding the completed turn. Furthermore, in the AMBER trajectories the hydrogen bonds at the terminus form after the hydrogen bonds at the turn, providing evidence for the hydrogen bond “zipping” process proposed by Muñoz *et al.* ^36,37^ and corroborated by Pande and Rokhsar ^12^.

**FIG. 4.**
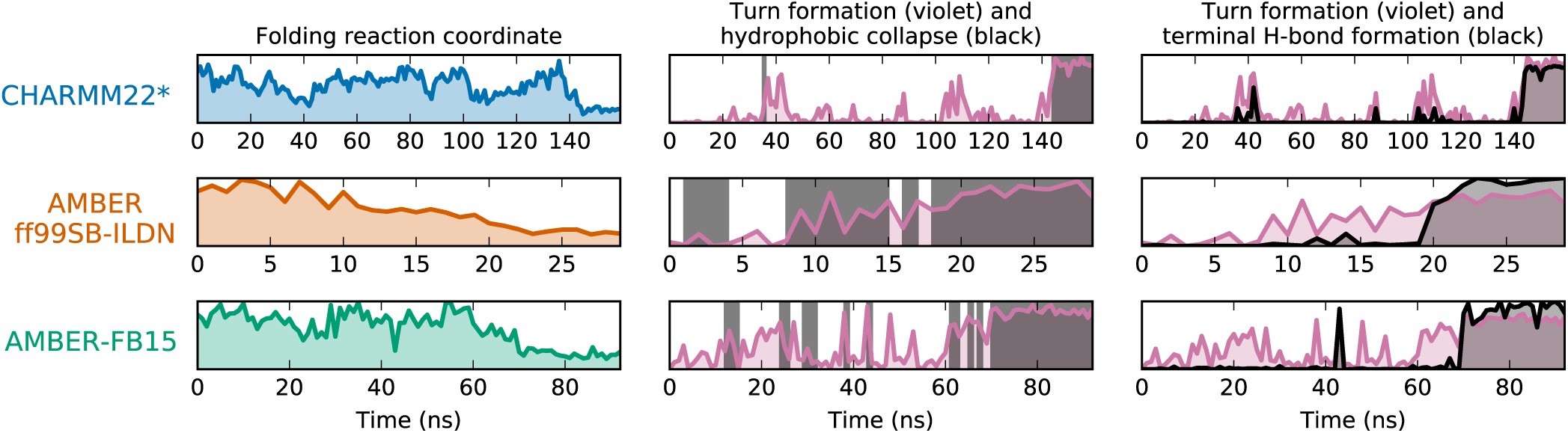
Representative folding trajectories reveal the mechanism of CLN025 folding. Each row represents a single trajectory that starts in an extended state and ends in the folded state. The left column shows the folding reaction coordinate (see Fig. 1), where low *y*-axis values correspond to the folded state. In the center and right columns, the formation of the three hydrogen bonds characterizing turn formation are shown in purple, where high values on the *y*-axis indicate the turn is formed. In the center panel, dark gray shading is used to show when the structure is collapsed. In the right panel, the formation of the three terminal hydrogen bonds is shown in dark gray, where high *y* values indicate that all three bonds are formed. In the CHARMM22* model, the collapse and formation of both turn and terminal hydrogen bonds occur nearly simultaneously. In the AMBER models, the appearance of collapsed structure precede the turn, and the turn hydrogen bonds form before the terminal hydrogen bonds.

Single trajectories are provided for interpretability. For this reason, visualization of at least 12 additional pathways for each force field are provided in the supplementary material. An example movie for each force field is also provided as a supplementary file.

### C. Rate-determining process

The original beta-hairpin formation theories also disagree on the rate-limiting step of the folding process. Muñoz *et al.* ^36,37^ hypothesize that the formation of the turn from the extended state determined the rate of beta-hairpin formation. Pande and Rokhsar ^12^ agree that the first step of the mechanism determined the rate, but in their mechanism the hydrophobic collapse preceded the turn and thus the collapse from the extended state characterized the rate-limiting step. The mechanism of Dinner, Lazaridis, and Karplus ^11^ identifies the rate-limiting step as the interconversion between collapsed conformations; i.e., the formation of the turn and native hydrogen bonds from a compact state.

In order to analyze the separate processes involved in beta-hairpin formation in our datasets, we constructed MSMs over two specific feature sets designed to characterize either hydrophobic collapse or turn formation. These sparse feature sets isolate the process of interest so that structures in the MD dataset are differentiated only by characteristics relevant to the appropriate process. Because the MSM timescales describe the timescales of conformational change, the longest timescale of each MSM corresponds to the relaxation time of each process of interest^55,77–79^. These values can be directly compared with process-specific experimental relaxation timescales^9^. In order to estimate the rate of hydrophobic collapse, the features selected were the radius of gyration of the two terminal hydrophobic residues (TYR1 and TYR10). To estimate the rate of turn formation, we calculated distances between the hydrogen bonded contacts in the turn region of the CLN025 crystal structure depicted in Fig. 3. The dihedral angles associated with these hydrogen bonds were also included, since it has been shown that only certain turn dihedrals can lead to the correct secondary structure^80^. We then constructed an optimized MSM for each feature set (see the supplementary material for feature descriptions, optimization protocol, and model validation). The per-model reaction coordinate and slowest relaxation timescale for these two processes are depicted in Fig. 5.

**FIG. 5.**
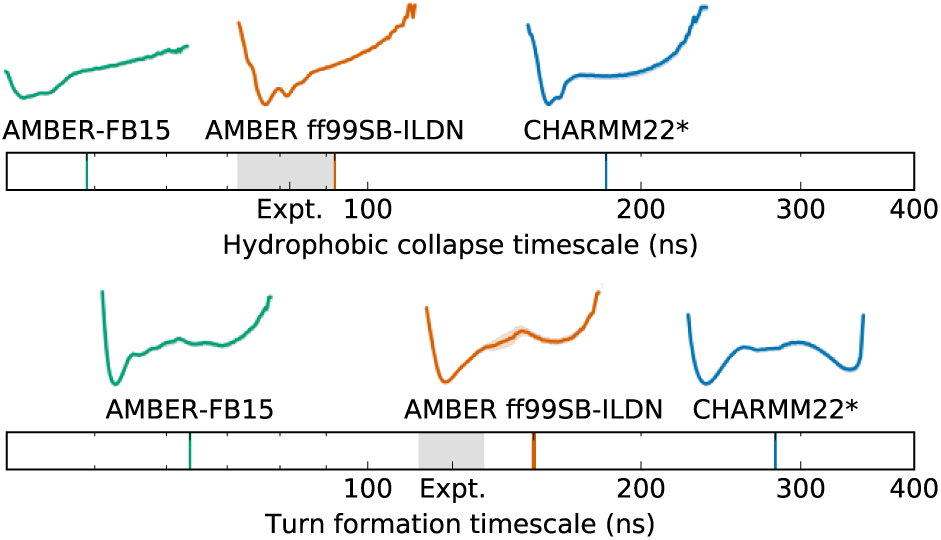
Isolating the processes involved in CLN025 folding reveals differences in their timescales and kinetics. The timescale plots show that all models calculate the turn formation to occur on a slower timescale than the hydrophobic collapse. The experimental values from Davis *et al.* ^9^ are represented by the gray shading. The reaction coordinate free energy plots above the timescale plots show that for all models hydrophobic collapse occurs downhill while turn formation occurs over a barrier. The rate-limiting step is the slower formation of the turn from the collapsed state. The ruggedness of the hydrophobic collapse landscapes indicates the existence of collapsed conformations in the denatured ensemble. Uncertainty in MSM timescale and experimental values are indicated by the thickness of the line. Shading represents the range of free energies based on 100 bootstrapped samples between the 5^th^ and 95^th^ percentiles.

First, we note that the relative ordering of timescales agrees with the experiments of Davis *et al.* ^9^. For all force field datasets, we observe a separation between the timescales corresponding to slower turn formation and faster hydrophobic collapse. Next, we note that the reaction coordinates corresponding to the hydrophobic collapse describe downhill pathways. In contrast, the reaction coordinates corresponding to turn formation feature a barrier between the turned and not turned conformations. From the relative ordering of timescales and the shape of the collapse and turn pathways, we agree with the conclusions of Davis *et al.* ^9^ and find that the rate-determining process for beta-hairpin formation is the formation of the turn from a pre-collapsed structure. This is also consistent with the rate-limiting step for beta-hairpin folding proposed by Dinner, Lazaridis, and Karplus ^11^.

## IV. DISCUSSION

In summary, our aggregated MD analysis suggests a beta-hairpin folding mechanism in which the extended state collapses into a hydrophobic cluster, followed by a slower process in which the hairpin turn forms over a barrier within the denatured collapsed state. The order of these conformational changes agree with the experimental conclusions reported by Davis *et al.* ^9^ for CLN025. Additionally, the resolution of MD simulations has allowed us to also model the formation of specific native state hydrogen bonds. We observe that the hydrogen bonds are formed by a “zipping” mechanism from the turn toward the terminus. Our findings demonstrate mixed agreement with the early theories of beta-hairpin formation; namely, our results support the “turn zipper” process of hydrogen bond formation^36,37^, the collapse-then-turn mechanism^12^, and the rate-determining process comprising rearrangement within a collapsed state^11^.

Performing this analysis simultaneously with datasets built from three different protein/water force field combinations demonstrates the force field dependence of CLN025 simulations at the experimental melting temperature. We find that simulations performed with CHARMM22* and AMBER ff99SB-ILDN yield an over-stabilized native state and unequal folding and unfolding rates, which indicates that a higher simulation temperature would be necessary to obtain melting behavior. In contrast, AMBER-FB15 simulations show behavior consistent with melting. Furthermore, while the folding mechanism can be determined using either AMBER dataset, the CHARMM22* dataset does not contain a compact denatured state at the simulated temperature, nor does it resolve the ordering of hydrogen bond formation via the “zipper” mechanism. We recommend that modelers who wish to use MD simulation to interrogate the denatured state ensemble of a protein and/or its role in the protein folding process choose a force field that accurately represents denatured state properties at the temperature of interest, and highlight that the AMBER-FB15 model yields behavior consistent with experiment at the simulated temperature. We anticipate that protein force fields that are accurate beyond the native state and sensitive to temperature dependence will enable further insight into larger and more complex protein systems.

Free, open source software implementing the methods used in this work is available in the OpenMM^59^, MDTraj^81^, MSMBuilder^82^, and Osprey^83^ packages available from http://openmm.org, http://mdtraj.org, and http://msmbuilder.org.

## SUPPLEMENTARY MATERIAL

Complete descriptions of all methods used in this work and additional data visualizations are available in the supplementary material. Example movies for CLN025 folding in each force field are provided as supplementary files. MSM objects compatible with the MSMBuilder software have been provided for all MSMs discussed in the main text. Details of these files and instructions for loading them can be found in the supplementary material.

## ACKNOWLEDGMENTS

The authors thank Lee-Ping Wang and Muneeb Sultan for helpful discussions and Jerelle Joseph for invaluable manuscript feedback. We acknowledge the National Institutes of Health under No. NIH R01-GM62868 for funding. We are also grateful to Caitlin Davis and Brian Dyer for discussions about their experimental results and to the Research Coordination Network: Protein Folding and Dynamics meeting funded by NSF MCB 1516959 for stimulating these discussions. We graciously acknowledge Folding@home donors who contributed to the AMBER force field simulations and D. E. Shaw Research for providing CHARMM22* simulation data. V.S.P. is a consultant & SAB member of Schrodinger, LLC and Globavir, sits on the Board of Directors of Apeel Inc, Freenome Inc, Omada Health, Patient Ping, Rigetti Computing, and is a General Partner at Andreessen Horowitz.

